# Immunomodulatory endothelial cells contribute to T cell recruitment and activation through antigen presentation on MHC class II

**DOI:** 10.1101/2025.07.17.665432

**Authors:** Matteo Cartura, Blerina Aliraj, Witold Szymański, Giulio Ferrario, Flávia Rezende, Beyza Güven, Tore Bleckwehl, Stefanie Dimmeler, Oliver J. Müller, Hanjoong Jo, Stefan Offermanns, Sikander Hayat, Ralf P. Brandes, Christian Münch, Johannes Graumann, Andreas Weigert, Ingrid Fleming, Mauro Siragusa

## Abstract

**Aims:** A subset of endothelial cells referred to as immunomodulatory endothelial cells (IMEC) has been proposed to regulate T cell responses in atherosclerosis, but their phenotype and function remain poorly understood. Here, we characterized the inflammation-induced emergence of IMEC and their crosstalk with T cells.

**Methods and Results:** An *in vitro* model to study IMEC was established and characterized using flow cytometry and proteomics. Single-cell transcriptome data from human atherosclerotic arteries as well as single cell transcriptome and endothelial cell-specific translatome data from a murine atherogenesis model were used to determine pathophysiological relevance. Immunopeptidomics was performed to detect antigen presentation. T cell chemotaxis, adhesion and activation were assessed through flow cytometry and microscopy. IMEC were induced by treating human endothelial cells with interleukin-1β, interferon-γ, and transforming growth factor-β2. These cells expressed lower levels of classical endothelial cell markers but expressed major histocompatibility complex (MHC) class II, proteins involved in antigen processing and presentation (CD83, CD80 and CD86) and pro-inflammatory cytokines as well as chemokines, including CXCL9. An endothelial cell subpopulation with similar immunomodulatory features was identified in a mouse model of accelerated atherogenesis as well as in human atheromas. Conditioned medium from IMEC enhanced the migration of peripheral blood mononuclear cells and induced T cell chemotaxis, the latter being partially inhibited by antagonizing CXCL9. Proteins related to glycosaminoglycan degradation were significantly downregulated in IMEC which was relevant inasmuch as the glycocalyx plays a key role in the establishment of chemokine gradients. Indeed, the accumulation of heparan sulfates in IMEC contributed to the adhesion of T cells. Notably, IMEC that had been exposed to monocyte lysates presented 627 peptide antigens on MHC class II and induced T cell activation.

**Conclusion:** Our data demonstrate the role of IMEC as non-professional antigen-presenting cells that potentially contribute to T cell-mediated immune responses in cardiovascular disease.

**Translational Perspective:** This study characterizes immunomodulatory endothelial cells (IMEC) as critical mediators of vascular inflammation through their capacity to process and present exogenous antigens and activate T cells. Induced by pro-atherogenic cytokines (IFN-γ, IL-1β, TGF-β2), IMEC upregulate MHC class II and costimulatory molecules, promote leukocyte chemotaxis, and enhance T cell adhesion through surface heparan sulfate. The identification of IMEC-like populations in both murine models and human atherosclerotic plaques indicates a conserved immunological function in atherogenesis. These findings position IMEC as novel, non-professional antigen-presenting cells and potential therapeutic targets to modulate vascular immune responses in atherosclerotic cardiovascular disease.

## Introduction

In inflammation, the presentation of antigens plays a key role in the amplification of the response and disease progression.^1^ Professional antigen presenting cells, such as dendritic cells, macrophages and B cells, take up cellular debris, which is then processed and presented on major histocompatibility complex (MHC) class II molecules to T cells. This process elicits T cell activation and proliferation.^2^ T cells, particularly Th1 cells, secrete pro-inflammatory cytokines like interferon (IFN)-γ that further activate other immune cells e.g. macrophages, to exacerbate inflammation.^3^

Endothelial dysfunction is the first step in the development of vascular disease and is characterized by the expression of adhesion molecules and the secretion of cytokines and chemokines by endothelial cells.^4^ The subsequent infiltration of innate and adaptive immune cells into the vessel wall occurs in the absence of exogenous pathogens and is therefore referred to as sterile inflammation.^5^ Recently, a subset of endothelial cells was identified in human atherosclerosis and murine arteries exposed to disturbed flow that expressed gene signatures indicative of phagocytosis or scavenging, antigen presentation and T cell recruitment.^6,7^ Given their potential role in modulating T cell responses, this subtype of endothelial cells was then referred to as immunomodulatory endothelial cells (IMEC).^8,9^ The emergence of the IMEC phenotype is most likely attributable to the pro-inflammatory microenvironment in the vessel wall, in particular to interleukin (IL)-1β, IFN-γ and transforming growth factor (TGF)-β2. Since the characterization of this endothelial cell subpopulation to-date has been limited to their transcriptional signature, the aim of this study was to characterize their phenotype and investigate possible mechanisms mediating crosstalk between IMEC and T cells.

## Methods

The authors declare that all supporting data are available within the article and its online supplementary files.

### Cell culture

#### Human endothelial cells

Human umbilical veins were obtained from local hospitals and endothelial cells were isolated and cultured as described previously ^10,11^ and used up to passage 4. Endothelial cells were cultured in ECGM2 medium (PromoCell, Heidelberg, Germany) containing 8% heat inactivated foetal bovine serum (FBS), gentamycin (25 µg/mL) non-essential amino acids (Thermo Fisher Scientific, Schwerte, Germany) and Na pyruvate (1 mmol/L, Sigma-Aldrich, Darmstadt, Germany). To induce the IMEC phenotype, 90% confluent endothelial cells were treated with a combination of IL-1β (50 ng/mL, Peprotech, Hamburg, Germany), IFN-γ (50 ng/mL, Peprotech, Hamburg, Germany), and TGF-β2 (5 ng/mL, Peprotech, Hamburg, Germany) for three consecutive days, replacing the medium daily.

#### Peripheral blood mononuclear cells (PB-MNC)

Human PB-MNC were isolated from buffy coats using a Ficoll-Paque (PAN-Biotech, Germany, Aidenbach) gradient with a density of 1.077 g/mL. The mononuclear cell layer was carefully collected after centrifugation at 440 x g for 35 minutes without break and acceleration. The collected cells were washed with PBS and treated with erythrocyte lysis buffer (PAN-Biotech, Germany, Aidenbach). Cells were then resuspended in RPMI-1640 medium (PromoCell, Heidelberg, Germany) supplemented with vitamins, non-essential amino acids, sodium pyruvate, streptomycin, and penicillin.

#### THP-1 and Jurkat cell lines

Human monocytic THP-1 cells (Cat. #TIB-202, ATCC, Germany, Wesel) and Jurkat E6-1 cells (Cat. #88042803, Merck, Darmstadt, Germany) were cultured in RPMI-1640 medium (PromoCell, Heidelberg, Germany) supplemented with 1 mmol/L sodium pyruvate and 10% FBS and 0.5% penicillin-streptomycin.

All cells used in this study tested negative to mycoplasma and were cultured in a 37 °C humidified incubator with 5% CO₂. The use of human material in this study complies with the principles outlined in the Declaration of Helsinki (World Medical Association, 2013), and the isolation of endothelial cells was approved in written form by the ethics committee of the Goethe-University.

### Animals

#### Endothelial cell-specific RiboTag mice

To generate endothelial cell-specific RiboTag mice (RiboTagEC), a model was developed by crossing endothelial cell-specific Cre-driver mice (Cdh5-CreERT2) with RiboTag mice (Rpl22HA/HA). The RiboTag model carries the ribosomal subunit Rpl22 allele with a floxed wild-type C-terminal exon, followed by an identical exon containing three hemagglutinin (HA) epitope copies inserted before the stop codon.^12^ The offspring were backcrossed to the C57BL/6 genetic background for at least 8–10 generations. Cre-mediated recombination was induced in 8–9-week-old mice via intraperitoneal tamoxifen injections (50 mg/kg per day in 50 µL Miglyol; Merck, Darmstadt, Germany) for five consecutive days. This enabled the expression of HA-tagged ribosomal subunit Rpl22 specifically in endothelial cells, integrating into actively translating polyribosomes. The tagged ribosomes were purified using an HA-specific monoclonal antibody, and ribosome-associated RNAs were sequenced and compared to total RNAs, enabling the analysis of translation changes in endothelial cell-specific transcripts.

#### AAV-PCSK9 generation and partial carotid artery ligation

An AAV serotype 8 vector for expression of the murine D377Y-PCSK9 cDNA (AAV-PCSK9) was produced using the two-plasmid-method by co-transfecting AAV/D377Y-mPCSK9 together with the helper plasmid pDP8 in HEK-293T cells using polyethylenimine (Merck, Darmstadt, Germany). AAV vectors were purified using iodixanol step gradients and titrated as described^13^. Accelerated endothelial activation and dysfunction was induced as described^14^. At day 0, mice were injected once with AAV-PCSK9 (10^11^ VG) via the tail vein and were fed a cholesterol rich diet (metabolizable energy 35 kJ% fat, containing 12.5 mg/kg cholesterol; EF PAIGEN, Ssniff, Soest, Germany). One week after initiation of the cholesterol rich diet, partial ligation of the left carotid artery was performed, as described.^15^ An injection of 0.1 mg/kg buprenorphine 30 minutes before surgery was used to alleviate surgical pain before, during and after awakening from anaesthesia. Metamizole (non-opioid analgesic, 1.4 mg/ml (350 mg/kg/day), total duration of 5 days) was administered via the drinking water to alleviate postoperative pain one day preoperatively and a further 3 days postoperatively. The animals were placed in an airtight anaesthesia box to induce anaesthesia, the oxygen-lsoflurane mixture was then flooded with 4% VE lsoflurane via an isoflurane vaporiser. After anaesthesia was induced, the animals were placed on a heated warming mat for the entire duration of the operation to prevent hypothermia, and the cornea of the eyes was protected from drying out with Bepanthen eye ointment. For the duration of the operation (approx. 15-20 min), the mice were sedated via a mask with approx. 1.5 - 3% VE lsofluran VE as required. On the day of organ collection, the animals were placed in an airtight anaesthesia box and the oxygen-isoflurane mixture was then flooded with >5% isoflurane via an isoflurane vaporiser. Animal death was confirmed by the complete cessation of breathing for at least one minute and the absence of paw reflex responses. All animal experiments were performed in accordance with the Directive 2010/63/EU of the European Parliament on the protection of animals used for scientific purposes and approved by the Federal Authority for Animal Research at the Regierungspräsidium Darmstadt (Hessen, Germany) under study protocols B2/2077.

Animal sample size was calculated using Software G*Power Version 3.1.9.7. No animals were excluded from any of the experiments. Animals were randomly assigned to the different groups and there was no source of confounders as all animals were handled similarly and were subjected to the same treatments and surgical procedures. Given the exploratory nature of the study, blinding was not applied.

### Flow cytometry

#### Human cells

Cells were harvested, washed and resuspended in PBS containing 0.5% BSA. Samples were blocked with 2% Fc Receptor Blocking Reagent (Miltenyi Biotec, Bergisch Gladbach, Germany) incubated with fluorochrome-coupled antibodies. Samples were acquired with a FACSymphony A5SE cytometer (BD Bioscience, Heidelberg, Germany) and the data was analyzed in FlowJo™ v10.9.0 Software (BD Life Sciences, Heidelberg, Germany). All primary antibodies were titrated to determine the optimal concentration. Compensation Beads (BD Bioscience, Heidelberg, Germany) were used for single-color compensation to create multicolor compensation matrices. For the gating strategy, fluorescence minus one control were applied. The instrument was controlled daily by calibrations with Cytometer Setup and Tracking beads (BD Bioscience, Heidelberg, Germany).

| Antigen | Dye | Dilution | Company | Catalog |
| --- | --- | --- | --- | --- |
| CD80 | BV711 | 1:50 | BD Bioscience, Heidelberg, Germany | 568227 |
| CD86 | FITC | 1:100 | Proteintech Hamburg, Germany | FITC-65/65 |
| HLA-DR | APC-Fire750 | 1:50 | BioLegend, Koblenz, Germany | 307658 |
| CD3 | BUV805 | 1:200 | BD Bioscience, Heidelberg, Germany | 612893 |
| CD45RA | BV711 | 1:200 | BD Bioscience, Heidelberg, Germany | 563733 |
| CD69 | BUV395 | 1:100 | BD Bioscience, Heidelberg, Germany | 564364 |
| CD45RO | AF700 | 1:100 | BD Bioscience, Heidelberg, Germany | 561136 |
| CD279 | FITC | 1:50 | BD Bioscience, Heidelberg, Germany | 557860 |
| CD4 | RB613 | 1:100 | BD Bioscience, Heidelberg, Germany | 571074 |
| CD197 | BV421 | 1:50 | BioLegend, Koblenz, Germany | 353208 |
| CD62L | BUV496 | 1:50 | BD Bioscience, Heidelberg, Germany | 750589 |
| TCRgd | BUV615 | 1:50 | BD Bioscience, Heidelberg, Germany | 751308 |
| CD127 | BB700 | 1:50 | BD Bioscience, Heidelberg, Germany | 566398 |
| CD8 | BV650 | 1:200 | BD Bioscience, Heidelberg, Germany | 563821 |
| CD25 | PE-Cy7 | 1:50 | BD Bioscience, Heidelberg, Germany | 557741 |
| HLA-DR | APC/Fire750 | 1:50 | BioLegend, Koblenz, Germany | 307658 |

### Immunofluorescence and microscopy

Cells were incubated with an antibody against MHC II (2 µg/ml, Cat. #NBP3-08644H, Novus Biologicals, Wiesbaden Nordenstadt, Germany) or perlecan (5 µg/ml, Cat. #13-4400, Thermofisher scientific, Darmstadt, Germany) at 37°C for 1 hour followed by incubation with an antibody against VE-Cadherin (5 µg/ml, Cat. #AF938, R&D Systems, Wiesbaden Nordenstadt, Germany). Cells were then washed with PBS and fixed in 4% paraformaldehyde (Carl Roth, Karlsruhe, Germany) at room temperature for 15 minutes. Blocking was performed with PBS containing 5% horse serum for 30 minutes at room temperature. Cells were then incubated with donkey anti-rabbit Alexa Fluor 555 (6.6 µg/ml, Cat. #A-21428, Thermo Fisher Scientific, Darmstadt, Germany) and donkey anti-goat Alexa Fluor 647 (6.6 µg/ml, Cat. #A-21244, Thermo Fisher Scientific, Darmstadt, Germany) in PBS for 1 hour at room temperature. Nuclei were stained using 4′,6-diamidino-2-phenylindole (DAPI) (0.1 µg/mL; AppliChem, Darmstadt, Germany) diluted in PBS. Finally, cells were overlaid with a mounting medium containing 50% (v/v) glycerol (Cat. #3783.1; Carl Roth, Karlsruhe, Germany) and dithiothreitol (DTT, 1 mol/L) (Cat. #A2948; AppliChem, Darmstadt, Germany) in PBS. Images were taken using a confocal microscope (STELLARIS 8 FALCON; Leica, Wetzlar, Germany) and LAS AF lite software (Leica). Bright field images of endothelial cells and IMEC grown on 8-well chamber slides (Ibidi, Martinsried, Germany) were acquired using confocal microscope (AXIO Vert.A1, Zeiss, EOS 700 D, Canon).

### Whole cell proteome analysis by mass spectrometry

Human endothelial cells (passage 4, 80% confluency) and IMEC were lysed in an S-trap-compatible buffer containing 5% SDS, 100 mmol/L triethylammonium bicarbonate buffer (TEAB), protease inhibitor cocktail (Roche EDTA-free) and supplemented with reducing (5 mmol/L tris(2-carboxyethyl)phosphine, TCEP) and alkylating (25 mmol/L Chloroacetamide, CAA) agents. The samples were incubated for 10 minutes at 95°C and sonicated with a Sonic Vibra Cell at 1s ON/1s OFF pulse for 30s at 40% amplitude to shear genomic DNA. After centrifugation at 10,000×g for 10 minutes at room temperature, the supernatant was transferred into fresh tubes and protein concentration was assessed using a BCA assay (Thermo Fisher, Darmstadt, Germany). Thirty µg of protein from each sample was used for S-trap sample preparation following the manufacturer’s protocol with few modifications. Briefly, samples (25 µL) were acidified adding 2.5 µL of phosphoric acid and subsequently washed with 175 µL of wash/binding buffer containing 90% methanol in 100 mmol/L TEAB. Samples were loaded into the S-trap spin column, spun at 4000xg for 1 minute to trap proteins and washed 5 times with washing buffer spinning the column at 4000xg for 30 seconds. After a final spinning cycle without any buffer, 20µ of digestion buffer (50 mmol/L TEAB) containing Trypsin (Promega, Walldorf, Germany) and Lys-C (Wako Chemicals, Neuss, Germany) were added, each in a 1:50 ratio overnight at 37°C. Peptides were eluted by three sequential steps of centrifugation at 4000xg for 1 minute with 40 µL of three different buffers; 50 mmol/L TEAB, 0.2% v/v Trifluoroacetic acid (TFA) and 50% v/v acetonitrile (ACN). The obtained peptides were dried and resuspended in 2% v/v ACN, 0,1% v/v TFA for C18 cleanup using a C18 plate (Affinisep). The plate was spun at 800xg for 1 minute, the C18 resin was activated with methanol, equilibrated with the same resuspension buffer. After that samples were loaded and washed 3 times with resuspension buffer. Elution was performed in 60% v/v ACN. Cleaned peptides were dried and resuspended in 30 µL of TMT labeling buffer (0.1 mol/L EPPS pH 8.2, 20% ACN). Fifteen µL of sample were mixed with TMTpro reagents (ThermoFisher Scientific, Darmstadt, Germany) in a 1:2.5 (w/w) ratio (2.5 µg TMT reagent per 1 µg peptide). Reactions were incubated for one hour at RT and after verification of labelling efficiency (>98%) and mixing ratios by LC-MS analysis of a test pool comprising 1/20th of each sample, labelling was quenched by addition of hydroxylamine to a final concentration of 0.5% and incubation at RT for 15min. Labelled peptides were pooled according to determined mixing ratios in order to achieve the same total peptide intensity in each sample and desalted by SepPak (tC18, 50 mg, Waters). Material was activated with methanol, followed by a wash with 80% acetonitrile (ACN) and equilibration with 5% ACN, 0.1% TFA. Samples were resuspended in 5% ACN, 0.1% TFA and loaded to resin material. Peptides were washed with 5% ACN, 0.1% TFA and eluted with 60% ACN and dried again. Peptides were fractionated using high-pH liquid-chromatography on a micro-flow HPLC (Dionex U3000 RSLC, Thermo Scientific, Darmstadt, Germany). Pooled and purified TMT labelled peptides resuspended in Solvent A (5 mmol/L ammonium-bicarbonate, 5% ACN) were separated on a C18 column (XSelect CSH, 1 mm x 150 mm, 3.5 µm particle size; Waters) using a multistep gradient from 3-60% Solvent B (5mmol/L ammonium-bicarbonate, 80% ACN) over 65 minutes at a flow rate of 30 µL/minute. Eluting peptides were collected every 43 seconds from minute 2 for 69 minutes into a total of 96 fractions, which were cross-concatenated into 24 fractions. Pooled fractions were dried in a vacuum concentrator and resuspended in 2% ACN, 0.1% TFA for LC-MS analysis. Tryptic peptides were analyzed on an Orbitrap Lumos coupled to an easy nLC 1200 (ThermoFisher Scientific, Darmstadt, Germany) using a 35 cm long, 75 µm ID fused-silica column packed in house with 1.9 µm C18 particles (Reprosil pur, Dr. Maisch), and kept at 50°C using an integrated column oven (Sonation). HPLC solvents consisted of 0.1% Formic acid in water (Buffer A) and 0.1% Formic acid, 80% acetonitrile in water (Buffer B). Assuming equal amounts in each fraction, 400 ng of peptides were eluted by a non-linear gradient from 7 to 40% buffer B over 90 minutes followed by a step-wise increase to 90% buffer B in 6 minutes, which was held for another 9 minutes. A synchronous precursor selection (SPS) multi-notch MS3 method was used in order to minimize ratio compression as previously described.^16^ Full scan MS spectra (350-1400 m/z) were acquired with a resolution of 120,000 at m/z 200, maximum injection time of 100 ms and AGC target value of 4 x 10^5^. The most intense precursors with a charge state between 2 and 6 per full scan were selected for fragmentation (“Top Speed” with a cycle time of 1.5 seconds) and isolated with a quadrupole isolation window of 0.7 Th. MS2 scans were performed in the Ion trap (Turbo) using a maximum injection time of 50 ms, AGC target value of 1.5 x 10^4^ and fragmented using CID with a normalized collision energy (NCE) of 35%. SPS-MS3 scans for quantification were performed on the 10 most intense MS2 fragment ions with an isolation window of 0.7 Th (MS) and 2 m/z (MS2). Ions were fragmented using HCD with an NCE of 50% (TMTpro) and analysed in the Orbitrap with a resolution of 50,000 at m/z 200, scan range of 100-500 m/z, AGC target value of 1.5 ×10^5^ and a maximum injection time of 86ms. Repeated sequencing of already acquired precursors was limited by setting a dynamic exclusion of 60 seconds and 7 ppm and advanced peak determination was deactivated. All spectra were acquired in centroid mode. Raw data was analyzed with Proteome Discoverer 2.4 (ThermoFisher Scientific, Darmstadt, Germany). Acquired MS2-spectra were searched against the human reference proteome (Taxonomy ID 10090) downloaded from UniProt (28-February-2024; “One Sequence Per Gene”, 20526 sequences) and a collection of common contaminants (244 entries from MaxQuant’s “contaminants.fasta”) using SequestHT, allowing a precursor mass tolerance of 7 ppm and a fragment mass tolerance of 0.5 Da after recalibration of mass errors using the Spectra RC-node applying default settings. In addition to standard dynamic (Oxidation on methionines and Met-loss at protein N-termini) and static (Carbamidomethylation on cysteins) modifications, TMT-labelling of N-termini and lysines were set as static modifications. False discovery rates were controlled using Percolator (< 1% FDR on PSM level). The PSM table was exported from Proteome Discoverer and subsequent data analysis was performed with Rstudio (2023.12.1 Build 402) with R version 4.3.2. PSM clean-up was based on using a QFeatures object-based analysis^17^. PSMs filtering was performed based on signal-to-noise above 10, at least 50% SPS-matches and a co-isolation below 75%. PSM-to-protein aggregation was performed according to robustSummary function from the MsCoreUtils package^18^. Only proteins quantified in all replicates in both experimental groups were used for statistical analysis. Differential expression analysis was performed using the limma package applying a q-value cut-off (<0.05) and fold-change threshold (log2ratio larger than +-0.58)^19^.

### Gene ontology pathway enrichment analysis

Differentially regulated proteins were used to perform gene ontology pathway enrichment analysis (STRING version 12.0 - ranked analysis - FDR ≤ 0.05).^20^

### Transcription factor analysis of immunomodulatory endothelial cells

Differentially expressed proteins identified by mass spectrometry were used as input for transcription factor (TF) enrichment analysis using ChEA3 ^21^, which identifies TF that may drive experimentally observed alterations in gene expression. The resulting list of candidate TF was then intersected with all proteins detected in IMEC compared to endothelial cells. All transcription factors that were not detected in the proteomic analysis were filtered out from the list of TF candidates. To evaluate their potential role in regulating the expression of genes involved in antigen processing and presentation, the top five upregulated TF candidates were further examined through cross-referencing with PubMed and GeneCards ^22^.

### Functional assays

#### T cell adhesion assay

Endothelial cells were starved for 1 hour in EBM containing 1% FBS and incubated for 30 minutes with solvent (DMSO, Sigma, Darmstadt, Germany) or surfen hydrate (20 µmol/L, Darmstadt, Germany). Surfen hydrate was dissolved in DMSO using glass or plastic materials pre-coated with FBS overnight. Forty thousand Jurkat cells were added in the presence of surfen hydrate or DMSO and allowed to adhere to endothelial cells for 2 hours. Non-adherent Jurkat cells were gently aspirated, and endothelial cells were washed with warm EBM containing 1% FBS. Cells were fixed with 4% paraformaldehyde for 10 minutes and washed with PBS. Images of fields with homogeneous distribution of endothelial cells were captured using an EVOS XL Core microscope (Invitrogen by Thermo Fisher Scientific) at 20x magnification. Quantification was performed using Fiji ImageJ with the “Count Particles” tool. Adhesion was expressed as number of Jurkat cells per image field.

#### Chemotaxis

Human endothelial cells or IMEC were exposed to basal RPMI media without supplements or FBS in the presence of a neutralizing antibody against CXCL9 (1 µg/ml, Cat. #MA5-23746, Thermo Fisher Scientific, Darmstadt, Germany) or isotype control IgG (1 µg/ml, Cat. #sc-2025, Santa Cruz Biotechnology, Heidelberg, Germany) for 4 hours. The conditioned culture media was collected and filtered to remove any potential cell debris (Filtropur S 0.2 µm, Sarstedt, Nümbrecht, Germany). Freshly isolated peripheral blood-derived mononuclear cells (PB-MNC) were resuspended in RPMI media and seeded into the upper chamber of a transwell insert (3 µm pore size, Sarstedt, Nümbrecht, Germany) at a density of 1×10⁶ cells. Inserts were then transferred to 24-well plates containing the conditioned culture media from human endothelial cells or IMEC. PB-MNC were allowed to migrate for 4 hours at 37°C in a humidified incubator with 5% CO₂, and migrated cells were quantified using the CASY cell counter.

### Single cell RNA-sequencing

#### Mouse atherogenesis

A study investigating how disturbed blood flow influences endothelial cell behaviour *in vivo* was used to assess the pathophysiological relevance of IMEC.^7^ Analyses were conducted at two days and two weeks after partial carotid ligation. The Seurat object containing pre-processed and normalized single-cell RNA sequencing data from this study was used to visualize the expression of transcripts for MHC class II and co-stimulatory molecules in IMEC and endothelial cells.

#### Human atheroma

A high-resolution map of human atherosclerotic plaques was generated by merging single-cell RNA sequencing and spatial transcriptomics data from 12 individuals.^6^ These single-cell RNA-sequencing data were analyzed using Seurat v5. Cells expressing either KDR or CDH5 and lacking expression of lymphatic markers (LYVE1, PROX1) were selected for further analysis of endothelial cells. The data was normalized, and 8,000 variable genes were identified. To account for donor-specific variations, batch correction was performed using harmony ^23^ and the resulting components were used for UMAP embedding. Clustering was evaluated across multiple resolutions based on marker gene expression, with resolution 0.4 selected for further analysis.

### RiboTag isolation and next generation sequencing

Murine carotid arteries were homogenized in lysis buffer (2-3% w/v) containing: Tris/HCl pH 7.5 (50 mmol/L), NaCl (150 mmol/L), MgCl_2_ (12 mmol/L), NP-40 (1%), DTT (1 mmol/L), EDTA-free protease inhibitor mix (AppliChem GmbH, Darmstadt, Germany), SUPERase-In RNase Inhibitor (200 U/mL; Thermo Fisher Scientific, Darmstadt, Germany), murine RNase inhibitor (500 U/mL; New England Biolabs, Frankfurt am Main, Germany), cycloheximide (100 µg/mL, Sigma-Aldrich, Darmstadt, Germany), TURBO DNase (25U/mL, Thermo Fisher Scientific, Darmstadt, Germany), and heparin (10 mg/mL, Thermo Fisher Scientific, Darmstadt, Germany) in UltraPure DNase/RNase-free distilled water (Thermo Fisher Scientific, Darmstadt, Germany). Lysates were centrifuged (16,000 g, 10 minutes, 4°C) and the supernatants passed through 70 µm pre-separation filters (Miltenyi Biotec, Bergisch Gladbach, Germany). A fraction (10%) of the total lysate volume was mixed with 9 volumes of QIAZOL (QIAGEN, Hilden, Germany) and frozen at −80°C. The remaining lysates were incubated with anti-HA antibody-conjugated magnetic beads (50 µL/sample, washed and resuspended in lysis buffer (Biozol, Eching, Germany) before being incubated overnight on an end-over-end rocker at 4°C. HA immunoprecipitates were washed 3 times with a freshly prepared high salt buffer containing: Tris/HCl pH 7.5 (50 mmol/L), NaCl (300 mmol/L), MgCl_2_ (12 mmol/L), NP-40 (0.5%), DTT (1 mmol/L), SUPERase-In RNase Inhibitor (200 U/mL), murine RNase inhibitor (500 U/mL) and cycloheximide (100 µg/mL). After the final wash, HA immunoprecipitates were suspended in 700 µL QIAZOL and immediately processed for RNA purification. The total input RNA (from the total lysates) as well as the ribosome-associated RNA (from the RiboTag immunoprecipitates) were purified using the miRNeasy Micro kit (QIAGEN, Hilden, Germany) according to the manufacturer’s instructions. RNA and library preparation integrity were verified with a LabChip Gx Touch 24 Nucleic Acid Analyzer (Perkin Elmer, Waltham, USA). Due to the low amount of RNA recovered from mouse carotid artery samples, only 10 ng of total RNA was used (SMART-Seq stranded kit, Takara Bio, Gennevilliers, France) according to the manufacturer’s instructions with standard RNA fragmentation (6 minutes, +85°C) and no further size selection. Sequencing was performed on the NextSeq500 instrument (Illumina) using v2 chemistry, resulting in 8-62M reads per library (average 25M) with 1×75 nt single-end setup.

### Bioinformatic analyses

#### QC/Trimming

Raw reads were assessed for quality, adapter content and duplication rates with FastQC 0.11.8 (Andrews S. 2010, FastQC: a quality control tool for high throughput sequence data: http://www.bioinformatics.babraham.ac.uk/projects/fastqc). Trimmomatic version 0.39 was employed to trim reads after a quality drop below a mean of Q15 in a window of 5 nucleotides for smORF analyses or a mean of Q20 in a window of 20 nucleotides for gene analyses.^24^ Only reads of at least 15 nucleotides were cleared for subsequent analyses.

#### Gene expression pipeline

Trimmed and filtered reads were aligned versus assembly version hg38 for human or mm10 for mouse data (Ensembl release 99 - 109) using STAR 2.6.1d or 2.7.10a to include multi-mapping reads with the parameters “—outFilterMismatchNoverLmax 0.1 --outFilterScoreMinOverLread 0.9 --outFilterMatchNminOverLread 0.9 --alignIntronMax 200000 --outFilterMultimapNmax 999”.^25^ The number of reads aligning to genes was counted with featureCounts 1.6.5 or 2.0.4 of the Subread package.^26^ Only reads mapping at least partially inside exons were admitted and aggregated per gene. Reads overlapping multiple genes or aligning to multiple regions were excluded. Differentially expressed genes were identified using DESeq2 version 1.18.1 or 1.36.^27^ Genes were classified to be significantly differentially expressed at average count > 5 with Benjamini-Hochberg corrected P value ≤ 0.05 and −0.585≤ Log2fold change ≥+0.585. The annotation was enriched with UniProt data (release 25.06.2019) based on Ensembl gene identifiers.^28^

### IMEC immunopeptidome

#### MHC class II immunoprecipitation

IMEC cultured in 15 cm plates were incubated overnight in the presence or absence of a cell lysate from 2.5×10^5^ THP-1 cells generated through three freeze-thaw cycles. IMEC were then washed three times with PBS and lysed in a lysis buffer containing NP-40 (1%), NaPPi (10 mmol/L), NaF (20 mmol/L), orthovanadate (2 mmol/L), okadaic acid (10 nmol/L), β-glycerophosphate (50 mmol/L), phenylmethylsulfonyl fluoride (230 µmol/L) and an EDTA-free protease inhibitor mix (AppliChem GmbH, Darmstadt, Germany). Lysates were incubated overnight with an antibody against MHC class II (2 µg antibody/1 mg of protein lysate, Cat. #NBP3-08644H, Novus Biologicals, Wiesbaden Nordenstadt, Germany) or isotype control IgG (2 µg antibody/1 mg of protein lysate, Cat. #NI01-100UG, Merck, Darmstadt, Germany) and protein A/G-sepharose 4B conjugate (30 µL slurry/1 mg protein lysate; Cat. #17061801; Cytiva, Dreieich, Germany). Immunoprecipitates were then washed three times with lysis buffer and peptides bound to MHC class II were eluted three times with 0.15% trifluoroacetic acid (Cat. #76-05-1, Merck, Darmstadt) at 37°C with agitation for 15 minutes.

#### Liquid chromatography and tandem mass spectrometry

Acidic eluates were loaded directly onto the C18 microspin columns and purified according to the manufacturer’s instructions (Macherey-Nagel, Düren, Germany), based on the original protocol.^29^ Purified peptides were first dried, then resuspended in 30 μL of 0.1% formic acid (FA, Thermo Scientific, Darmstadt, Germany), and 0.01% dodecyl-β-D-maltoside (DDM, Roth, Karlsruhe, Germany). Peptide concentration was estimated using the Pierce Fluorimetric Peptide Assay, and sample volumes were adjusted to achieve equal concentrations. Peptides were analyzed by liquid chromatography–tandem mass spectrometry (LC-MS/MS) carried out on a Bruker Daltonics timsTOF Ultra instrument connected to a Bruker Daltonics nanoElute instrument. Approximately 30 ng of peptides were loaded onto a C18 precolumn (Thermo Trap Cartridge, 5 mm; μ-Precolumn™ Cartridge / PepMap™ C18, Thermo Scientific, Darmstadt, Germany) and then eluted in backflush mode with a gradient from 95% solvent A (0.1% formic acid in water) and 5% solvent B (99.9% acetonitrile and 0.1% formic acid) to 15% solvent B over 5 min, then to 45% solvent B over an additional 25 min, using a reverse-phase high-performance liquid chromatography (HPLC) separation column (PepSep Ultra, C18, 1.5 μm, 75 μm × 25 cm, Bruker Daltonics, Bremen, Germany) at a flow rate of 300 nL/min. The column temperature was set to 50 °C. The outlet of the analytical column was coupled to the MS instrument via a CaptiveSpray 20 μm emitter. Data were acquired using a data-dependent acquisition (DDA) paradigm with a default method provided by Bruker (PS_dda_norm_0.5s.m). Briefly, spectra were acquired with fixed resolution of 45,000 and a mass range from 100 to 1700 m/z for the precursor ion spectra and a 1/k0 range from 0.7 to1.45 V s/cm2 with a 100 ms ramp time for ion mobility. Detailed method settings can be extracted from the deposited measurement files. Peptide spectrum matching and label-free quantification were subsequently performed using MaxQuant (Version 2.5.1.0)^30^ against the Human UniProt database (20,429 entries, November 2024). Digestion was set to unspecific, with the minimal peptide length set to 8 and the maximal length to 40 amino acids. Methionine oxidation and N-terminal acetylation were set as variable modifications. The full list of settings may be found in the “mqpar.xml” file uploaded to the ProteomeXchange repository. Downstream data processing and statistical analysis were carried out by the limma-based^19^ Autonomics package developed in-house (DOI: 10.18129/B9.bioc.autonomics; version 1.15.145). Proteins with a q-value of <0.01 were included for further analysis. Median-centered intensity values were used for quantification and missing values were imputed. Max Quant initially identified 1835 protein groups. All intensities that contained only 1 precursor per sample, were replaced with NA for that particular sample. After dropping proteins without replication (within a subgroup), and filtering out proteins with fewer than 2 peptides identified, 1,762 protein groups were retained for further analysis. 488 protein groups had systematic NAs [missing completely in some subgroups but detected in others (for at least half of the samples)], 208 protein groups had random NAs (missing in some samples, but unrelated to subgroup affiliation) and 34 protein groups had no NAs. Systematic NAs were imputed by random numbers drawn from a normal distribution with a width of 0.3 and a downshift of 1.6 standard deviations from the mean sample-wise, with the SD and distribution calculated for each group separately. Imputation is indicated in the final result tables.

#### MHC class II binder selection

For each MS-identified peptide, log_2_ fold changes were calculated for the following comparisons: (i) MHC class II vs. control IgG (basal), (ii) MHC class II + THP-1 vs. control IgG, and (iii) MHC class II + THP-1 vs. MHC class II. The first two comparisons were used to filter out peptides identified in the control IgG immunoprecipitates and to retain peptide significantly enriched in the MHC class II immunoprecipitates. In the third comparison, only peptides significantly enriched against the respective IgG controls in the first and second comparisons were retained. Peptides that failed to meet these criteria were categorized as false positives and excluded. To ensure that only high-confidence peptides were selected for further analysis, only peptides with an Andromeda score ≥ 15.326 were included. This cut-off score was chosen because it was the Andromeda score of the peptides derived from CD74, which was one of the most upregulated proteins in IMEC compared to endothelial cells in the independent proteomic analysis presented in this study. Peptides were further filtered based on the average reported size for MHC class II binders (15 to 25 amino acids).^31^

### Immunogenicity predictions

The immunogenicity of MHC class II peptide binders identified by immunopeptidomics was predicted using the CD4 T cell immunogenicity prediction tool from Immune Epitope Database.^32^ The sequence of each binder was compiled into a FASTA file and analysed using a method that combines MHC binding (7-allele method) and immunogenicity predictions with a percentile threshold of combined score 50, with low values indicating high capacity of being recognized by the T cell receptor. The sequences of tetanus and diphtheria toxoids were also included in the analysis as positive controls.^33,34^

### T cell activation

Endothelial cells or IMEC were treated with THP-1 protein lysate overnight, as described above. Cells were washed three times and freshly isolated PB-MNC were seeded on top at a density of 100,000 cells per 1 cm². After 72 hours, the cells were collected and resuspended in cold PBS with 0.5% BSA for flow cytometry analysis. Different T cell subpopulations were identified by the following markers: CD4^+^ T cells (CD45+, CD3+, CD4+, CD25-, CD127+/-), γδ T cells (CD45+, CD3+, CD4-, CD8-, gdTCR+), Tregs (CD45+, CD3+, CD4+, CD25+CD12low) and CD8+ T cells (CD45+, CD3+, CD8+).

### Statistics

Results are presented as mean ± SEM. GraphPad Prism software (v. 9 to 10.4.1) was used to assess statistical significance. All data passed the Shapiro-Wilk normality test. Differences between two groups were compared by two-tailed unpaired t-test. Differences between three or more groups were compared by one-way ANOVA followed by the Tukey’s multiple comparisons test. Experiments in which the effects of two variables were tested were analyzed by 2 way ANOVA followed by the Tukey’s multiple comparisons test. Differences were considered statistically significant when P < 0.05. Only exact significant P values are reported.

### Data availability

All mass spectrometry proteomics data have been deposited to the ProteomeXchange Consortium ^35^ via the PRIDE or MassIVE (https://massive.ucsd.edu/, MassIVE-ID: MSV000097582, dataset license: CC0 1.0 Universal).partner repositories^36^ with the dataset identifier PXD062928 (IMEC proteome) and PXD062803 (IMEC immunopeptidome).

NGS data have been deposited to the Gene Expression Omnibus with the accession number GSE195453.

## Results

### Stimulation of human endothelial cells with IL-1β, IFN-γ and TGF-β2 induces the IMEC phenotype

IL-1β, IFN-γ, and TGF-β2 are key cytokines that shape the immune landscape of atherosclerotic plaques.^37–39^ Therefore, endothelial cells were treated with these cytokines, individually and in combination, for three days. A cocktail containing all three cytokines induced the surface expression of MHC class II molecules as well as the co-stimulatory molecules CD80 and CD86 (**Fig. 1A**), which are required for antigen presentation and effective T cell activation.^40,41^ IFN-γ, on its own, was sufficient to induce the expression of MHC class II and CD86, but not CD80. These changes in protein expression were concomitant with marked alterations in endothelial cell morphology; cells stimulated with the cytokine cocktail lost their cobblestone morphology became elongated, and demonstrated a disrupted pattern of VE-cadherin expression (**Fig. 1B**).

**Figure 1.**
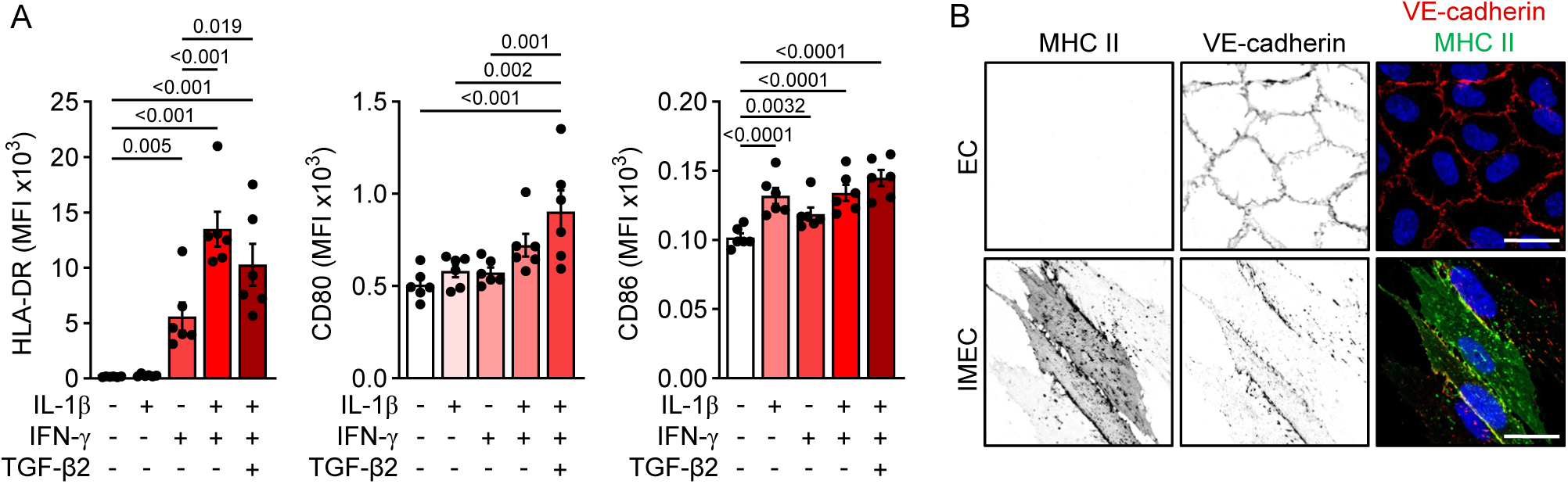
Endothelial cell phenotype induced by IL-1β, IFN-γ and TGF-β2. **A**, Expression of HLA-DR, CD80, CD86 on the surface of human endothelial cells after stimulation with IL-1β (50 ng/mL), IFN-γ (50 ng/mL) and TGF-β2 (5 mg/mL) for three days; n=6 independent cell batches (one-way ANOVA and Tukey multiple comparisons test). **B**, Representative images showing MHC class II and VE-cadherin in human endothelial cells (EC) and IMEC; bar = 20 µm. Similar images were obtained in 5 independent cell batches.

### IMEC express proteins involved in T cell chemotaxis and antigen presentation

To determine the molecular landscape of the IMEC phenotype, the IMEC proteome was compared with that of endothelial cells (from matched cell batches). Approximately 3000 proteins were differentially regulated, with 1,595 being down- and 1,435 up-regulated (−log_10_Pvalue > 1.3, −0.585 ≤ log_2_(IMEC/EC) ≥ 0.585) in IMEC (**Fig. 2A** and **Table S1**). Cytokine treatment led to the downregulation of endothelial markers such as CD31, CD34, vWF, EMCN, NOS3, DLL4, TIE1 and TEK compared to human endothelial cells (**Fig. 2B**). Interestingly, CDH5 was only slightly downregulated, KDR was not altered, and FLT1 was slightly upregulated. The most highly upregulated proteins were associated with the regulation of the immune system; including CXCL9, IFI27, ISG15, CD38, and CD83. Pathway enrichment analysis highlighted the expected upregulation of proteins related to IFN signaling as well as cytokine signaling, chemokines and their receptors, antigen presentation (MHC I), MHC class II protein complex and phagocytic vesicle membranes (**Fig. 2C**-**D**). The proteins most downregulated by induction of the IMEC phenotype were implicated in cell communication and adhesion (GJA5, CD34), receptor modulation and signaling (RAMP2) and immune system regulation (IL33, CCL14). Pathway enrichment analysis of all downregulated proteins revealed that chromatin remodelling (chromosome and chromatin organization, histone modification), as well as metabolic pathways (oxidative phosphorylation, DNA metabolic processes, and ceramide catabolism), were negatively affected in IMEC (**Fig. 2E**).

**Figure 2.**
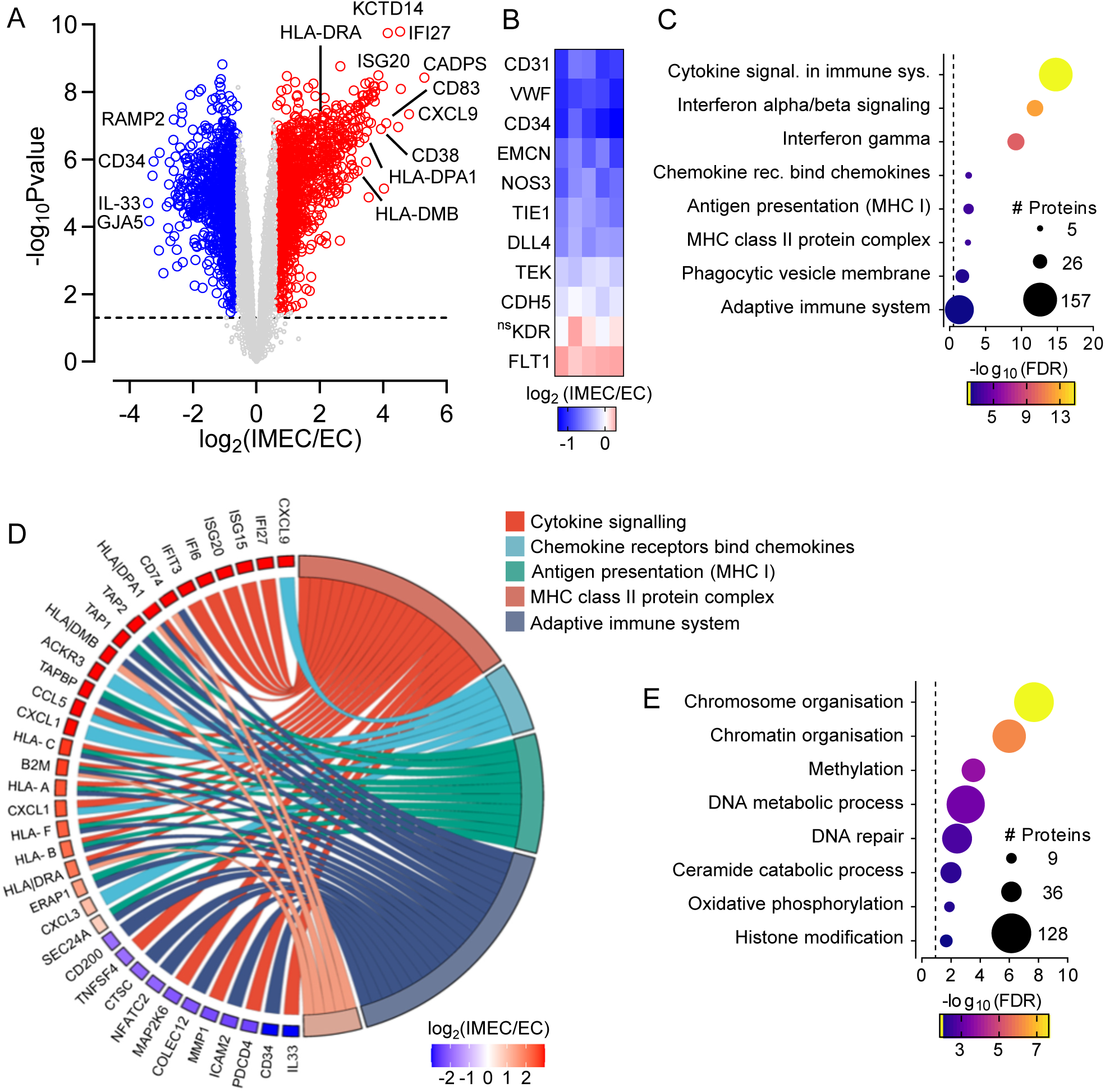
The IMEC proteome. **A**, Volcano plot showing differentially expressed proteins in IMEC compared to human endothelial cells (EC); n=5 independent cell batches. The dashed line marks the significance threshold (P value: 0.05). **B**, Heatmap showing the expression of endothelial cell markers in IMEC versus EC. **C**, Pathway enrichment analysis (STRING) of significantly upregulated proteins in IMEC compared to human endothelial cells. The dashed line marks the significance threshold (FDR: 0.05). **D**, Chord diagram displaying the top 5 upregulated and downregulated proteins in the top 5 enriched pathways, with proteins sorted based on log_2_ (IMEC/EC). **E**, Pathway enrichment analysis (STRING) of significantly downregulated proteins in IMEC compared to human endothelial cells. The dashed line marks the significance threshold (FDR: 0.05).

All of the differentially expressed proteins identified by mass spectrometry were used as input for transcription factor (TF) enrichment analysis using ChEA3,^21^ which identifies TFs that may drive experimentally observed alterations in gene expression (**Table S2**). The resulting list of candidates was then intersected with all proteins detected in IMEC compared to endothelial cells to filter out the TFs not detected in the IMEC proteome. This identified PLSCR1, SP100, numerous members of the IRF and NFκ-B families, SOX18, ELK3 and ERG as the major contributors to proteomic changes. Perhaps more importantly, the analysis also revealed that the 5 most upregulated TFs in the proteomic analysis i.e. SP110, MAFF, POU2F2, MAFK and STAT2 directly modulated proteins involved in antigen presentation and T cell activation (**Table 1** and **Table S2**). This fits well with the previous reports of SP110 and STAT2 being induced by IFN-γ^42,43^, and MAFF and MAFK being regulated by IL-1β and TGF-β2, respectively.^44,45^

**Table 1.**
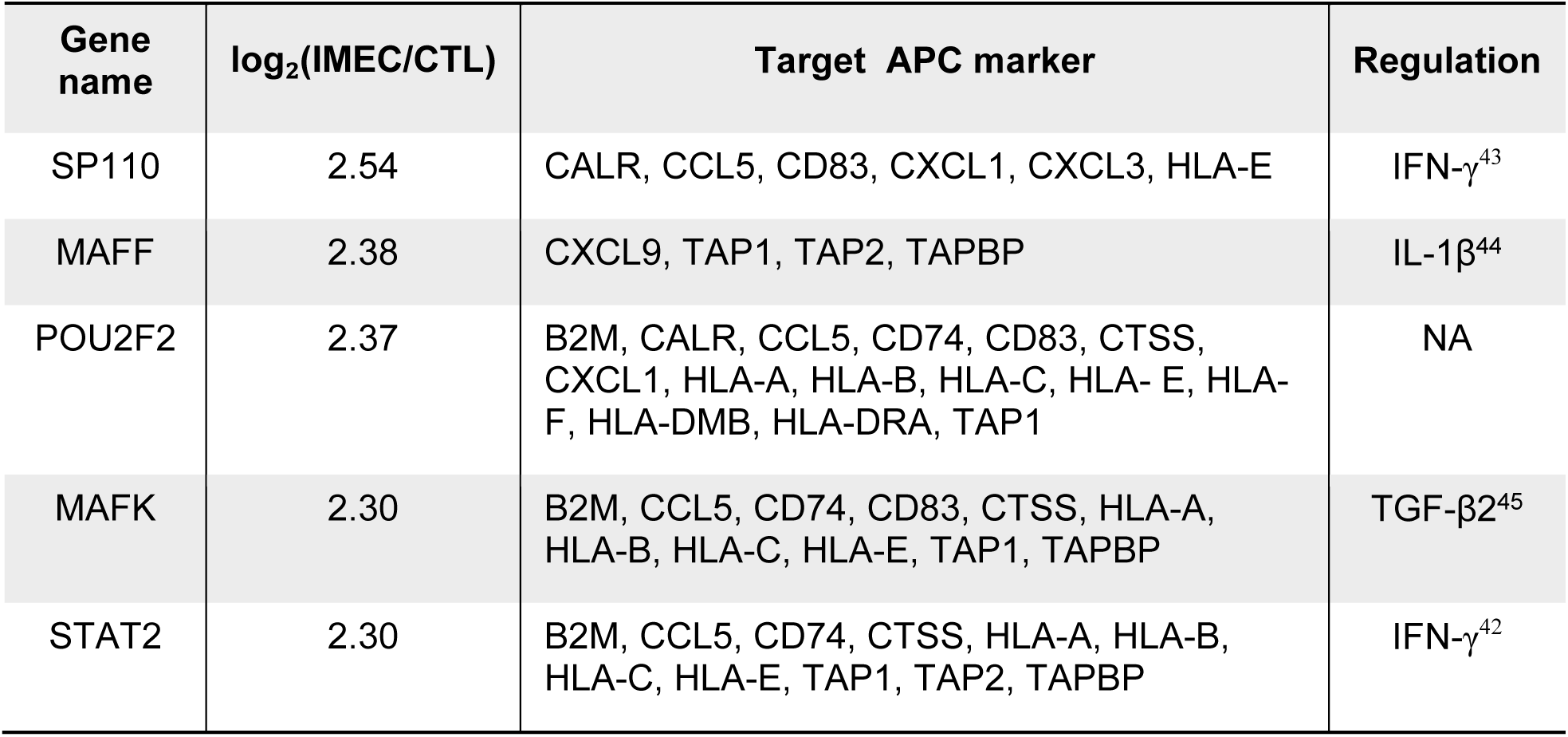
Top 5 transcription factors upregulated in IMEC and modulating expression of APC markers.

### IMECs recruit leukocytes via CXCL9 and promote T cell adhesion to heparan sulfate

Deeper interrogation of the proteome revealed that key enzymes involved in glycosaminoglycan degradation (KEGG database), were markedly downregulated in the IMEC proteome (**Fig. 3A**). Indeed, levels of five of the nine proteins involved in the degradation of heparan sulfate i.e., IDUA, SGSH, NAGLU, GUSB, and GNS, were clearly downregulated in IMEC, hinting that the IMEC surface is richer in heparan sulfate proteoglycans than the surface of endothelial cells. Fitting with this, levels of perlecan were higher in IMEC (**Fig. 3B**). Next, we assessed the impact of heparan sulfate accumulation on T cell adhesion to IMEC. A significantly larger number of T cells adhered to IMEC than endothelial cells, an effect that was strongly inhibited by treatment with the small HS antagonist; surfen hydrate (**Fig. 3C**).

**Figure 3.**
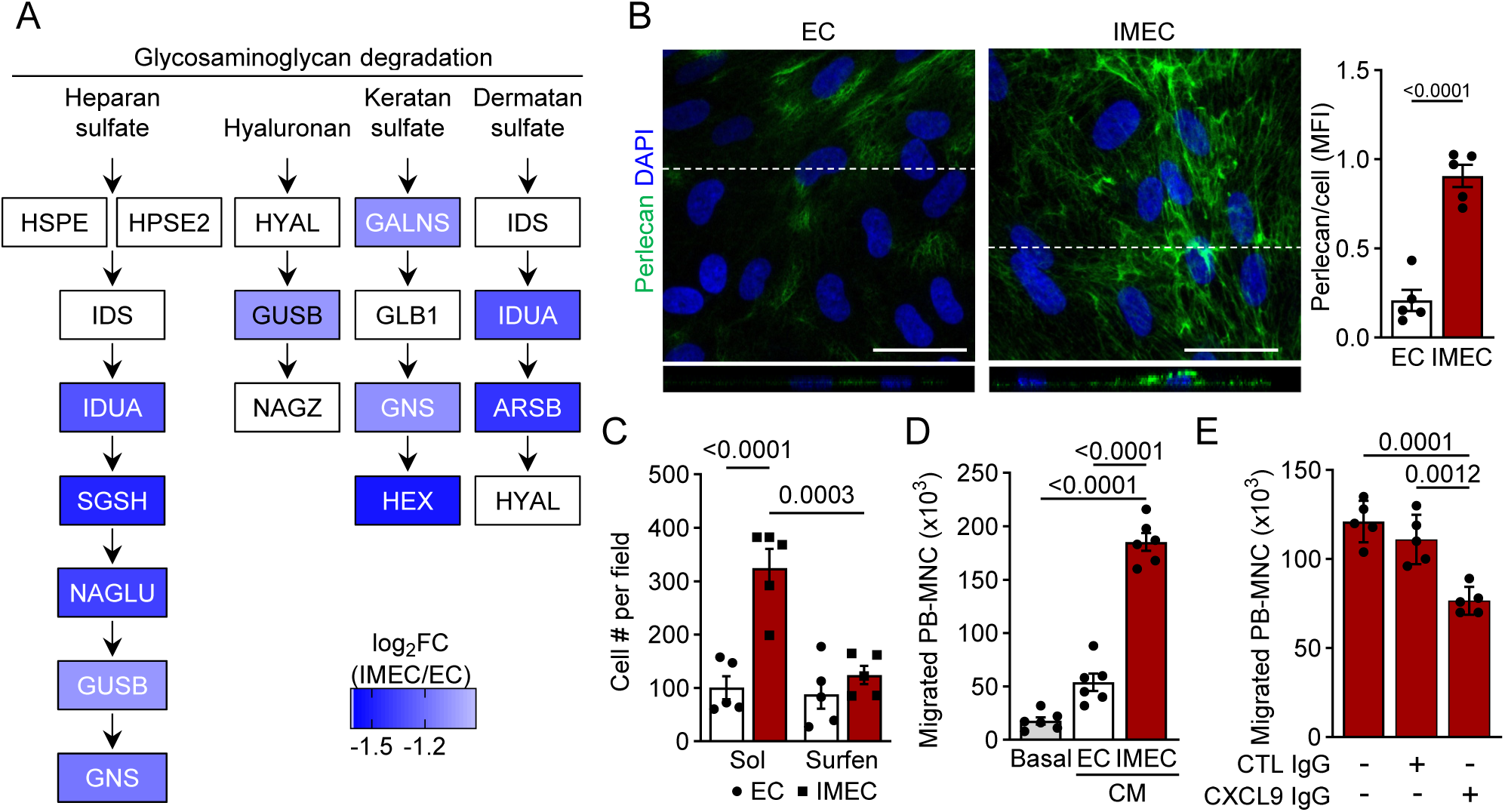
CXCL9-mediated leukocyte recruitment and T cell adhesion to heparan sulfate. **A,** Differentially expressed proteins in the glycosaminoglycan degradation pathway (KEGG) in IMEC compared to human endothelial cells (EC) from proteome analysis as in Fig. 2A**. B,** Perlecan levels in EC or IMEC. Scale bar=20 μm; n=5 independent cell batches (unpaired Student’s t-test). **C,** Adhesion of Jurkat cells to EC or IMEC treated with surfen hydrate or solvent (Sol); n=5 independent cell batches (two-way ANOVA and Tukey multiple comparisons test). **D,** PB-MNC migration toward conditioned media (CM) from EC, IMEC or RPMI (basal) for 4 hours; n=6 independent cell batches (one-way ANOVA and Tukey multiple comparisons test). **E,** PB-MNC migration toward IMEC CM in the presence of control IgG or a neutralizing antibody against human CXCL9; n=5 independent cell batches (one-way ANOVA and Tukey multiple comparisons test).

The endothelial glycocalyx is essential for the formation of chemokine gradients,^46^ which are in turn essential for guiding immune cells to sites of inflammation.^47^ To determine whether IMEC could establish such a strong gradient, a chemotaxis assay was performed to determine whether IMEC-derived factors could promote leukocytes recruitment. Compared to conditioned medium (CM) from endothelial cells, the IMEC supernatant induced significantly higher migration of peripheral blood-derived mononuclear cells (PB-MNC) (**Fig. 3D**). Whole cell proteome analysis indicated that IMEC modulate T cell recruitment via CXCL9. Consistent with this observation, incubation of PB-MNC with a neutralizing antibody against CXCL9 markedly reduced migration in response to IMEC-CM (**Fig. 3E**).

### Detection of IMEC in murine and human atherosclerosis

A subpopulation of endothelial cells characterized by an immune-like gene signature was previously identified in murine carotid arteries exposed to disturbed flow.^7^ To evaluate how closely the *in vitro* differentiated IMEC resembled the *in vivo* immune-like subpopulation, we interrogated the murine scRNA-sequencing dataset for the key antigen presenting cell markers that characterized the human IMEC proteome.

Two days after partial carotid artery ligation the expression of transcripts related to antigen processing and presentation i.e., MHC class II, CD74, CD80, CD83 and CD86 was higher than in the non-ligated right carotid artery (**Fig. 4A**). This difference was even more pronounced after 2 weeks. To confirm the occurrence of endothelial cells with antigen-presenting potential in an independent dataset, endothelial cell-specific RiboTag mice were made hypercholesterolemic by overexpression of PCSK9 and a high fat diet before partial carotid artery ligation was performed. Seven days after ligation, which represents the peak inflammatory response,^48^ the arteries were harvested and actively translated genes were identified by RiboTag RNA sequencing. Also in these experiments, MHC class II, CD74 and the co-stimulatory molecule Cd86 were all increased in carotid artery endothelial cells from ligated carotid arteries (**Fig. 4B** and **Table S3**).

**Figure 4.**
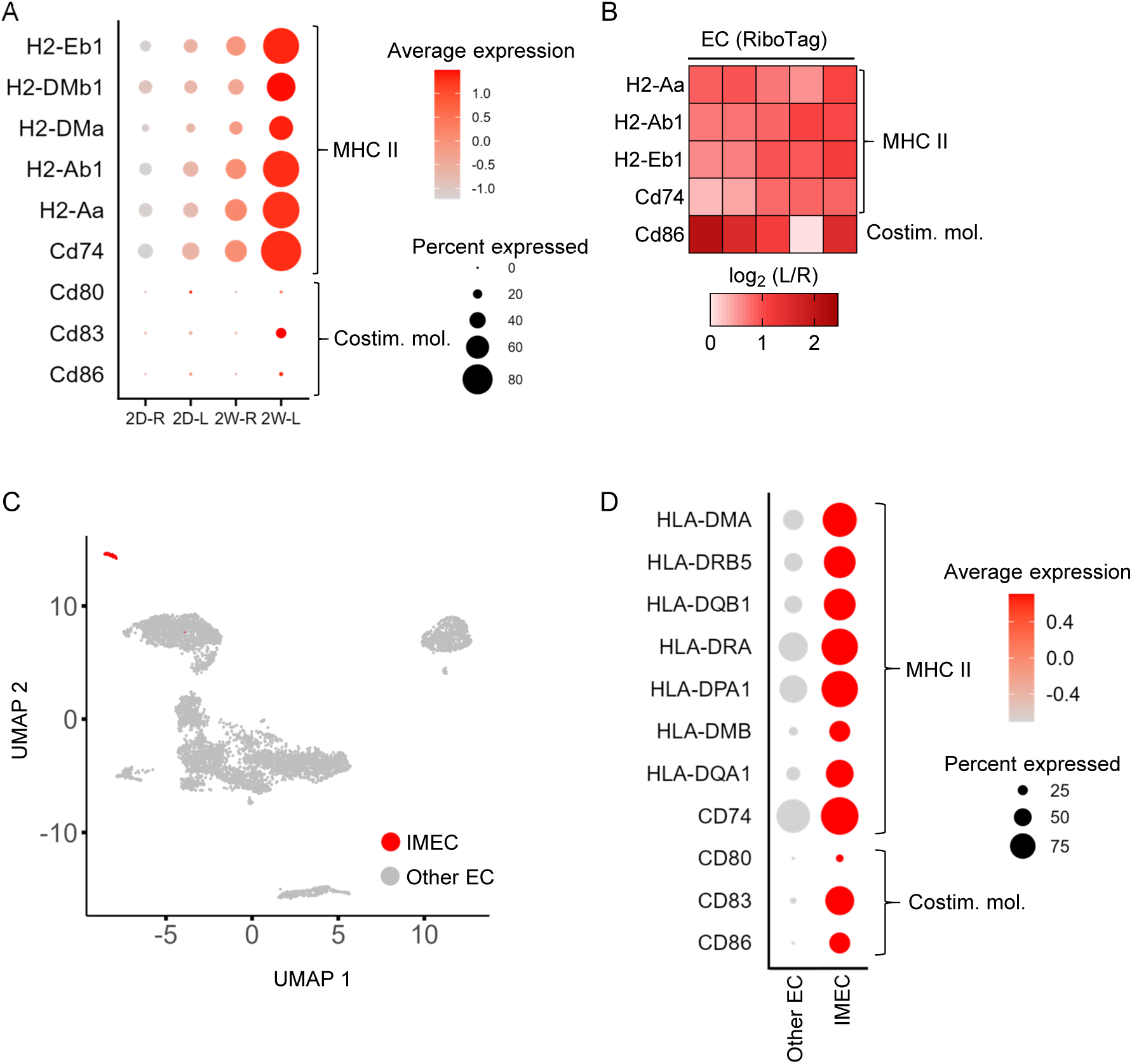
Identification of IMEC in murine arteries exposed to atherogenic disturbed flow and in human atherosclerosis. **A,** Expression of transcripts for MHC class II and co-stimulatory molecules in endothelial cells from ligated left (L) or right (R) murine carotid arteries 2 days (2D) or 2 weeks (2W) after partial carotid artery ligation. Bubble size indicates the percentage of cells expressing a given transcript and the colour scale represents average expression level. Carotid arteries were pooled from 10 mice to obtain single cells. **B,** Expression (RiboTag RNA sequencing) of MHC class II and Cd86 in the endothelium of ligated left (L) compared to control right (R) carotid arteries from hypercholesterolemic endothelial cell-specific RiboTag mice (EC RiboTag) 7 days after partial carotid artery ligation; n=5 mice. **C,** UMAP showing endothelial cell clusters identified by the expression of CDH5 and/or KDR and lack of Prox1 and Lyve1. The IMEC population is highlighted in red. n=12 patients with atherosclerosis. **D**, Expression of transcripts for MHC class II and co-stimulatory molecules in IMEC compared to other endothelial cells in human atheromas. Bubble size indicates the percentage of cells expressing a given transcript and the colour scale represents average expression level.

To identify an IMEC-like population in the human situation, we analyzed scRNA-seq data in the human atherosclerosis atlas.^6^ As the IMEC proteome lost classical endothelial cell markers but retained the expression of CDH5 and KDR (see **Fig. 2B**), we restricted our analyses to cells expressing CDH5 and/or KDR but lacking the lymphatic markers (LYVE1, PROX1). Among the endothelial cell clusters identified, a small IMEC-like population was detected (**Fig. 4C**). This population expressed MHC class II molecules and CD74, as well as the costimulatory molecules CD83, CD86 and CD80 (**Fig. 4D**).

### IMEC present antigens on MHC class II and activate T cells

To assess the ability of IMEC to process and present antigens on MHC class II molecules, the cells were treated with culture medium or a lysate derived from monocytes as a source of protein. MHC class II molecules were then immunoprecipitated from IMEC (**Fig. 5A**) and the MHC II-bound peptides were identified by mass spectrometry. Only peptides enriched in the MHC class II pulldowns (versus control IgG) that were 15 to 25 amino acids in length were considered to be *bona fide* MHC class II binders (**Table S4**). Under basal conditions, we detected 25 peptides originating from 7 proteins, including ITG5A and ECE1, which are enriched in endothelial cells, and CD74, involved in the stabilization of peptide-free MHC class II molecules (**Fig. 5B and D**). In contrast, IMEC treated with the monocyte lysate presented 565 peptides derived from 200 proteins (**Fig. 5C-D**). The top-presented peptide originated from NAP1L1, a protein preferentially expressed in monocytes, but many of the other peptides detected exhibited low cell type specificity. To predict MHC binding and generate immunogenicity scores, the peptide antigens identified were used as input in the CD4^+^ T cell immunogenicity prediction tool available in the Immune Epitope Database analysis resource.^32^ This identified a large number of immunogenic peptides with a combined score similar to that of immunogenic peptides derived from tetanus and diphtheria toxins (**Table S5**).

**Figure 5.**
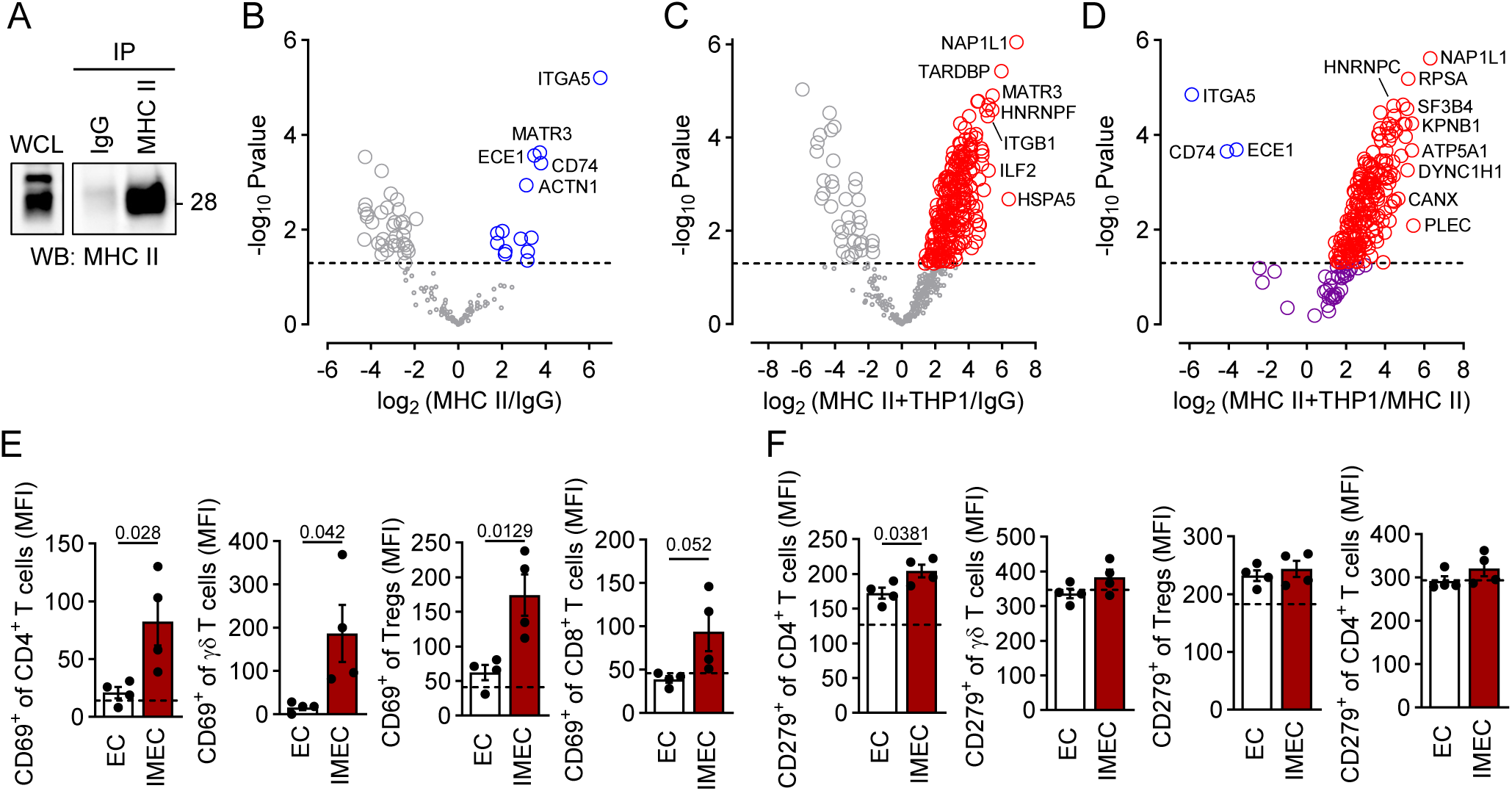
Antigen presentation and T cell activation by IMEC. **A,** Representative immunoblot showing the efficiency of the immunoprecipitation (IP) of MHC class II from whole cell lysate (WCL) of human IMEC. IgG was included as a control. Similar results were obtained in 3 independent cell batches. **B,** Volcano plot showing mass spectrometry-detected peptides enriched in MHC class II immunoprecipitates compared to control IgG from human IMEC under basal conditions. **C**, Volcano plot showing mass spectrometry-detected peptides enriched in MHC class II immunoprecipitates compared to control IgG from human IMEC treated with monocyte lysate. **D**, Volcano plot showing mass spectrometry-detected peptides enriched in MHC class II immunoprecipitates from IMEC treated with monocyte lysate versus IMEC under basal conditions. B-D, n=3 independent cell batches/condition. The dashed lines mark the significance threshold (Pvalue: 0.05). **E,** Expression of CD69 in CD4+ T cells (CD45+, CD3+, CD4+, CD25-,CD127+/-); γδ T cells (CD45+, CD3+, CD4-, CD8-, gdTCR+), Tregs (CD45+, CD3+, CD4+, CD25+CD12low) and CD8 + T cells (CD45+, CD3+, CD8+) three days after incubation of PB-MNC with endothelial cells or IMEC. Dashed lines indicate basal activation level of CD69 in PB-MNC alone. n=4 independent cell batches (unpaired Student’s t-test). **F,** Expression of CD279 in CD4^+^ T cells, γδ T cells, Tregs, and CD8 ^+^ T cells three days after incubation of PB-MNC with endothelial cells or IMEC, both treated with THP-1 protein lysate. Dashed lines indicate basal activation level of CD279 in PB-MNC alone. only n=4 independent cell batches (unpaired Student’s t-test).

T cell responses differ significantly based on the nature of the antigen, the T cell subset involved and the surrounding microenvironment.^49^ To determine whether IMEC were able to induce T cell activation, they were incubated with monocyte lysate to induce antigen presentation before being co-cultured with PB-MNC for 72 hours. Co-culture with IMEC elicited T cell activation evidenced by the increased expression of the early activation marker CD69 in CD4^+^ T cells, γδ T cells, regulatory T cells (Tregs) as well as CD8^+^ T cells (**Fig. 5E**). The expression of the late T cell activation marker CD279 increased after co-culture with IMEC only in CD4^+^ T cells (**Fig. 5F**). These results demonstrate that IMEC can function as non-professional antigen-presenting cells capable of initiating a broad T cell response. However, the restricted induction of CD279 in CD4⁺ T cells implies that IMEC may preferentially support sustained activation, differentiation or regulatory programming in this subset.

## Discussion

MHC class II is generally assumed to represent a class of inflammatory cells but to be absent from non-immune cells such as the vascular endothelium. The intriguing identification of a population of immunomodulatory endothelial cells in human atherosclerosis,^6^ as well as a subpopulation of endothelial cells with an immune-like gene signature in murine carotid arteries exposed to disturbed flow,^7^ has challenged this general assumption. In this study, it was possible to differentiate quiescent human endothelial cells into an IMEC phenotype and to demonstrate that IMEC present antigenic peptides via MHC class II molecules that are able to activate T cells.

In line with previous reports demonstrating transcriptional upregulation of genes linked to antigen processing and presentation in endothelial cells under inflammatory conditions,^6–9,50^ we found that a cocktail of cytokines implicated in atherogenesis i.e., IL-1β, IFN-γ, and TGF-β2 induced the IMEC phenotype. Consistent with its well-established role in modulating MHC class II gene expression in other cells,^50^ IFN-γ alone induced a robust surface expression of MHC class II. However, only the combination of IL-1β, IFN-γ, and TGF-β2 resulted in the concomitant surface expression of MHC class II as well as of the essential co-stimulatory molecules, CD80, CD86 and CD83. The latter protein was one of the most upregulated proteins in the *in vitro* differentiated IMEC and is relevant inasmuch as CD83 is highly expressed in mature dendritic cells and is thought to define cells that have acquired full antigen-presenting capability.^51,52^ The phenotypic switch from endothelial cell to IMEC was also associated with a decrease in the expression of proteins involved in chromatin organization, remodeling, and methylation, which may reflect an increase in chromatin accessibility and the establishment of a more permissive epigenetic landscape for transcriptional reprogramming. The transcription factor profile was also altered in IMEC with SP110, MAFF, POU2F2 (Oct-2), MAFK and STAT2 being the five most upregulated transcription factors. These observations fit well with reports that, with the exception of POU2F2, the transcription factors are directly modulated by IL-1β, IFN-γ or TGF-β2.^42–45^ POU2F2 levels are, however, controlled by NF-κB, which is activated by both IL-1β and IFN-γ,^53^ and is particularly interesting as it is implicated in the regulation of the Class II Transactivator (CIITA), which is a master regulator of MHC class II gene expression.^54^ Other transcription factors were negatively regulated by differentiation to IMEC; including SOX18 and the ETS transcription factors ELK3 and ERG, which are all well-established regulators of endothelial cell specification.^55–58^

The phenotypic switch from endothelial cell to IMEC was also associated with a number of changes that included the acquisition of an elongated shape with numerous protrusions. IMEC partially lost their endothelial cell identity, as indicated by the significant downregulation of endothelial cell markers. CDH5 is a classical endothelial cell marker and an important functional constituent of tight junctions and although protein levels were not markedly decreased, CDH5 patterning and localization on the cell surface was clearly disrupted, which is highly indicative of disrupted barrier function. Other downregulated proteins affecting intercellular communication included the gap junctional protein connexin 40 (GJA5),^59^ which was the second most significantly decreased protein in IMEC relative to quiescent endothelial cells. Among the proteins most significantly downregulated in IMEC was RAMP2, which is a co-receptor that enables the calcitonin receptor-like receptor to function as an adrenomedullin receptor.^60^ This observation is relevant as endothelial cells exposed to disturbed flow release adrenomedullin, which acts via its receptor to suppress proinflammatory signaling pathways. Indeed, the endothelial cell-specific deletion of adrenomedullin or its receptor in murine models leads to enhanced atherosclerosis.^61^ Our observations imply that IMEC are likely to be adrenomedullin-insensitive and that the loss of this key response could also contribute to atherogenesis. The proteomic analysis also identified an unexpected pathway that was markedly affected by differentiation to IMEC i.e., glycosaminoglycan degradation. Indeed, five out of nine enzymes in this pathway were clearly decreased in IMEC versus quiescent endothelial cells. This would be expected to have consequences on the expression of different glycosaminoglycans and it was possible to demonstrate that IMEC express higher levels of perlecan. This class of proteins is important as they are constituents of the endothelial cell glycocalyx which plays a critical role in the sensing of hemodynamic stimuli,^62^ as well as in leukocyte chemotaxis and adhesion by facilitating the formation of chemokine gradients.^46^

Another characteristic of the IMEC phenotype was the downregulation of the vasoprotective and anti-inflammatory mediator IL-33,^63,64^ and the parallel increase in the expression of pro-inflammatory mediators such as IL-1β, IFN-γ and IL-32, as well as the chemokines CXCL1, CXCL3 and CXCL9. While CXCL1 and CXCL are involved in the recruitment of neutrophils, CXCL9 is the major chemoattractant for T cells and was the most upregulated chemokine and the second most upregulated protein in IMECs. The functional relevance of this finding was demonstrated experimentally as IMEC recruited PB-MNC in a CXCL-9-dependent manner. Demonstrating such interactions *in situ* is a challenge but a recent study that combined spatial transcriptomics and scRNA-seq data identified an IMEC-like population of endothelial cells in close proximity to T cells in the *vasa vasorum* of human atherosclerotic plaques.^6^ These data highlight the pathophysiological relevance of IMEC *in vivo* and provide a hint about the location where the crosstalk between IMEC and T cells is likely to occur.

Having established that IMEC express key molecules involved in antigen processing and presentation, and are capable of attracting T cells, the next step was to determine whether IMEC can alter T cell function. Specifically, we assessed the ability of IMEC to internalize monocyte-derived proteins, process them and present the derived peptides on MHC class II molecules to activate T cells. Our immunopeptidomic analyses using IMEC revealed that, in the absence of monocyte lysate, MHC class II molecules were predominantly loaded with peptides derived from the invariant chain protein CD74. This is not surprising as this protein plays a central role in MHC class II antigen processing by stabilizing peptide-free α/β heterodimers and directing their trafficking from the endoplasmic reticulum to endosomal/lysosomal compartments for subsequent peptide loading.^65^ After exposure to a monocyte lysate, IMEC MHC class II molecules bound 580 peptides, some of which were predicted to have high immunogenicity and T cell-activating potential. This prediction was supported by functional assays in which robust T cell activation was demonstrated. While early T cell activation, as indicated by CD69 expression, was observed in CD4⁺, CD8⁺, γδ T cells and Tregs exposed to IMEC, the expression of the late activation marker; CD279, was selectively expressed by CD4⁺ cells. CD279 plays a dual role in mediating both the prolonged activation and the initiation of regulatory or exhaustion pathways,^66,67^ and its selective expression may reflect a mechanism by which IMEC contribute to the fine-tuning of adaptive immunity. This subset-specific upregulation may indicate that IMEC preferentially supports the sustained activation or functional modulation of CD4⁺ cells, possibly balancing activation with immune tolerance. It is important to note that although the T cell activation assays using a monocyte cell lysate confirmed the immunogenicity of monocyte-derived peptides, the *in vivo* repertoire of immunogenic peptides in atherosclerosis is likely to be very different. In the disease setting, antigens originate primarily from vascular smooth muscle cells, macrophages and foam cells that die during lesion progression and thus would be expected to contribute to a distinct antigenic landscape that my affect different T cell subsets.

Taken together, we demonstrated the role of IMEC as functional non-professional antigen-presenting cells and identified key molecular mechanisms mediating their crosstalk with T cells. As it was possible to detect an IMEC-like population in a mouse model of atherogenesis and in human atherosclerosis, these findings are relevant for vascular disease.

## Acknowledgements

The authors are indebted to Isabel Winter and Sandrine Ngaha for expert technical assistance.

